# Maize inbreds show allelic variation for diel transcription patterns

**DOI:** 10.1101/2024.12.16.628400

**Authors:** Joseph L. Gage, M. Cinta Romay, Edward S. Buckler

## Abstract

Circadian entrainment and external cues can cause gene transcript abundance to oscillate throughout the day, and these patterns of diel transcript oscillation vary across genes and plant species. Less is known about within-species allelic variation for diel patterns of transcript oscillation, or about how regulatory sequence variation influences diel transcription patterns. In this study, we evaluated diel transcript abundance for 24 diverse maize inbred lines. We observed extensive natural variation in diel transcription patterns, with two-fold variation in the number of genes that oscillate over the course of the day. A convolutional neural network trained to predict oscillation from promoter sequence identified sequences previously reported as binding motifs for known circadian clock genes in other plant systems. Genes showing diel transcription patterns that cosegregate with promoter sequence haplotypes are enriched for associations with photoperiod sensitivity and may have been indirect targets of selection as maize was adapted to longer day lengths at higher latitudes. These findings support the idea that cis-regulatory sequence variation influences patterns of gene expression, which in turn can have effects on phenotypic plasticity and local adaptation.

## Introduction

Because most plants rely on sunlight for energy, they need to entrain their biological processes to the rhythms of the sun. Plant growth, development, physiology, metabolism, and immunity are all governed by diel cues and the internal circadian clock(Lu *et al*. 2017), including temporal patterns of transcription for about one third of genes(Ferrari *et al*. 2019). The number and types of genes showing diel patterns of transcription vary across species in Archaeplastida, but core components of the circadian clock are conserved across green plants(Ferrari *et al*. 2019; Michael 2022; Petersen *et al*. 2022).

Despite high conservation of core circadian regulators, plant species have adapted to environments that vary wildly in their diurnal conditions, showing differences in aspects such as day length, light availability, temperature fluctuation, and pathogen pressure. These same conserved circadian regulators underpin molecular processes for circadian-regulated metabolic products that differentiate species from each other. Some of the diversity in form and function related to circadian processes can be attributed to genetic diversity in circadian genes, as previously shown in Arabidopsis(Michael *et al*. 2003) soybean(Wang *et al*. 2022) and maize(Hung *et al*. 2012b), but rewiring of gene interactions by evolution of transcription factor binding sites (TFBS) or chromatin accessibility are other, less explored ways that circadian regulation may have diversified among plant species.

Transcription factors (TFs) tend to evolve slowly and be conserved at both the sequence and functional level(Lambert *et al*. 2018; Tu *et al*. 2020). Their binding sites, on the order of ∼10bp in length(Stewart *et al*. 2012), experience much faster rates of evolution(Stone and Wray 2001; Weirauch and Hughes 2010; Lambert *et al*. 2018). The fact that TFBS evolve much faster than their cognate TFs makes TFBS evolution a strong possible explanation for the observed variability in diel gene expression patterns between species, despite a conserved core circadian network. In addition to between-species variability, differential activity of TFBSs may contribute to within-species diversity for diel gene expression. To date, many studies of diel or circadian gene expression patterns in plants have focused on a representative variety, mutant lines, or comparisons across species. The extent of within-species, natural allelic variation for diel gene expression patterns is less well characterized.

Maize is morphologically and genetically diverse, and has adapted to a wide range of environments worldwide, including broad latitudinal adaptation. Maize was originally domesticated in the Balsas valley of Mexico under short day lengths of less than 13 hours(Matsuoka *et al*. 2002; Piperno *et al*. 2009). Prior to European colonization, people living in the Americas moved maize to latitudes spanning from Chile to Canada(Sauer and Weatherwax 1955; Crawford *et al*. 2006), necessitating selection for growth and reproduction under long day conditions with up to 16 hour days. Short-day-adapted material grown in long-day environments will exhibit delayed flowering or potentially not flower at all; temperate-adapted maize varieties must be photoperiod-insensitive in order to produce grain.

Early flowering is a highly polygenic trait critical for adaptation to temperate environments at higher latitudes with longer days(Troyer and Hendrickson 2007; Buckler *et al*. 2009), whereas photoperiod sensitivity, while still polygenic, appears to be governed by fewer loci(Hung *et al*. 2012b). Genetic variation in the maize circadian clock gene *ZmCCT*, homolog of *Ghd7* in rice and partially similar to CONSTANS in Arabidopsis, has been shown to contribute to photoperiod sensitivity, but adaptation to long days is polygenic and *ZmCCT* accounts for 9% of phenotypic variance for photoperiod sensitivity in the maize Nested Association Mapping population(Hung *et al*. 2012b). Adaptation to long days likely required selection on a standing variation at a large number of loci(Swarts *et al*. 2017), but few of those have been rigorously identified or characterized, and the mechanisms by which selection altered flowering time in maize are not well known.

Here we describe natural allelic variation for diel patterns of transcript abundance among 24 maize inbred lines. We find that transcription factor binding motifs previously identified in Arabidopsis are predictive of whether or not a gene shows cyclical diel patterns of transcript abundance in maize. In addition, promoter region haplotypes co-segregate with significantly different rhythms of diel transcript abundance for hundreds of genes among our 24 inbreds. Those genes are shown to be enriched for GWAS hits for flowering and photoperiod sensitivity traits, and in some populations, enriched for signals of selection to higher latitudes. We posit that during movement to higher latitudes, people selecting for adaptation to longer days modified circadian regulation networks.

## Results and Discussion

### Natural variation for diel transcriptome rhythmicity

Twenty-four diverse inbred lines, a subset of the parents of the maize NAM population(Yu *et al*. 2008; Gage *et al*. 2020), were evaluated for transcript abundance every two hours for twenty-four hours (Supplemental Data 1). To reduce reference bias, sequencing data from each inbred were aligned to that inbred’s cognate genome assembly(Hufford *et al*. 2021). Each annotated gene in each inbred was evaluated for rhythmicity using diffCircadian(Ding *et al*. 2021), which uses a likelihood-based method to test whether diel transcriptomic data fit a sinusoidal function. This method is less likely to identify significant diel patterns of expression that are not sinusoidal, such as impulses, step changes, or sawtooth patterns.

We found variability between inbred lines for the number of genes which display significant diel rhythmicity (likelihood ratio test p < 0.01), ranging from 3,345 (Ms71) to 6,730 (Il14H) (Figure 1A, Supplemental File 1). The two genotypes with the highest number of rhythmic genes, P39 and Il14H, are both sweet corn lines which are genetically distinct from the remainder of the NAM parent lines, adapted to northern latitudes, and flower quickly. The number of cycling transcripts in a given inbred was weakly but not significantly correlated with photoperiod sensitivity, flowering time, maturity at sample collection, leaf sampled, or sequencing depth (Supplemental Figure 1), supporting the possibility that this pattern may be due to genetic differences in diel gene regulation and not simply varying levels of adaptation to the northern field site (Aurora, NY) where this study was performed.

**Figure 1:**
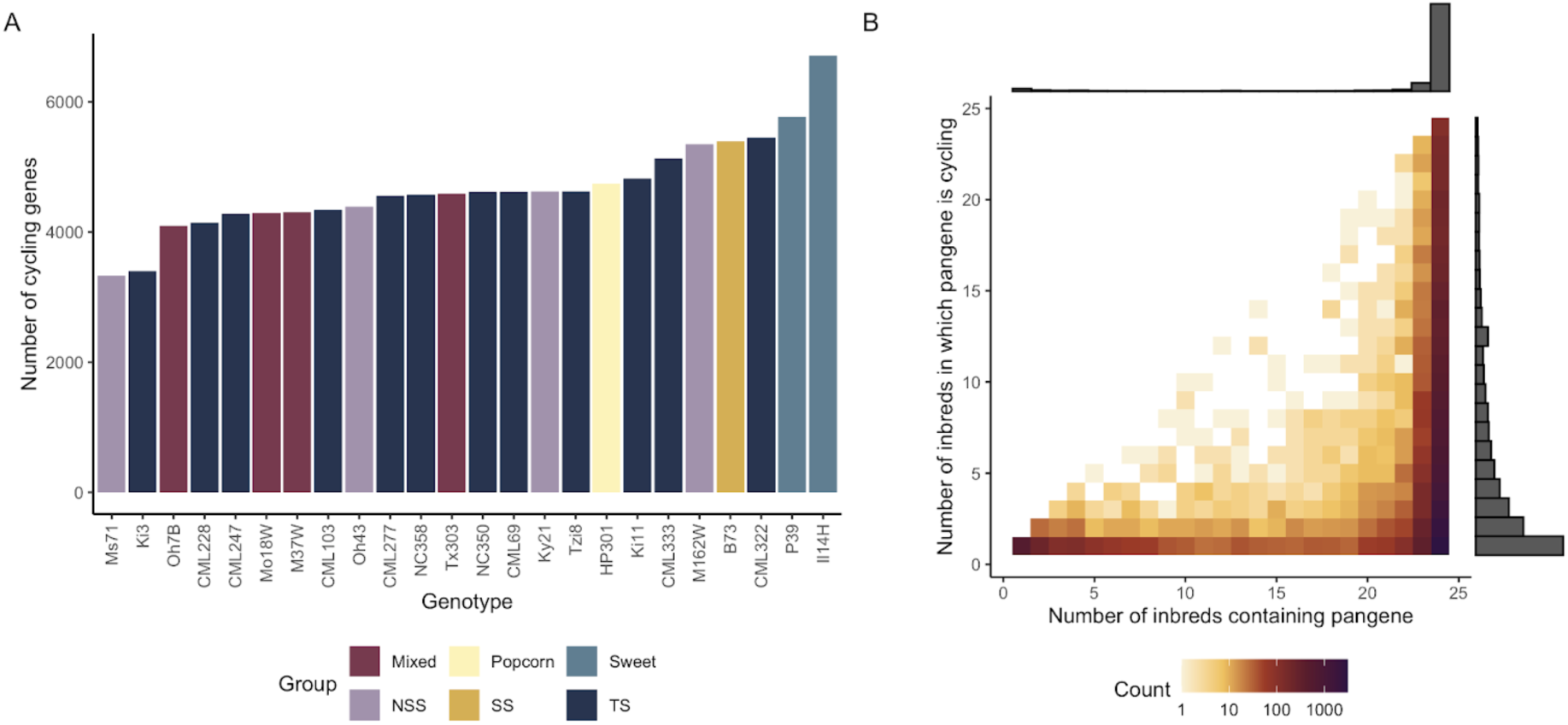
(A) Number of genes showing significant (p<0.01) sinusoidal diel patterns of transcript abundance in each inbred maize line. (B) Pangenes that show diel cycling are most likely to be present in all 24 inbred lines, but are often cycling only in some inbreds.

Using a pangenome(Hufford *et al*. 2021) to establish homologous pan-genes, we found that pan-genes which display diel rhythmicity in at least one inbred are frequently core genes (ie, present in all inbreds), but are not necessarily rhythmic in all inbreds (Figure 1B). This finding indicates that standing variation for rhythmicity exists in otherwise conserved genes and could have been selected upon during breeding and adaptation to new environments.

### Predicting rhythmicity from promoter sequences

Given that there is natural variation in diel transcription patterns, both among pan-genes and between alleles of pan-genes, we trained a convolutional neural network (CNN) to predict whether a given gene is likely to show a cyclical pattern of diel transcription (Supplemental Data 2). Based on previous characterization of the plant circadian clock, we know that diel transcription is subject to temporal feedback loops involving a number of transcription factors (TFs), including CCA1, LHY, PRRs, and ELFs(Sanchez and Kay 2016). We predicted that the presence, absence, and interactions between the binding motifs for these TFs might be identified from the convolutional filters of our model.

Among the motifs identified by our model (Supplemental Figure 2, Supplemental Data 3) was a strong representation of the Evening Element AAATATCT (Nagel *et al*. 2015). We also identified motifs related to other circadian TFs previously characterized in Arabidopsis, including the CONSTANS-responsive element CCACA (CORE2) (Gnesutta *et al*. 2017); the TOC1 TCP Binding Site (TBS) binding site motif GGCCC(Gendron *et al*. 2012); the TOC1 GA element(Gendron *et al*. 2012); the E2F/DP binding motif GGCGG(Vandepoele *et al*. 2005) which controls elements of the cell cycle, a process coupled to the circadian clock(Torii *et al*. 2022); and the Telo-box motif AAACCCT(Spensley *et al*. 2009; Gardiner *et al*. 2021). Though our model was able to identify known TF binding motifs, most of its predictive power is from predicting across pan-genes; it is limited in its ability to predict differences between pan-gene alleles.

These results reinforce conserved aspects of the plant circadian clock(Khan *et al*. 2010; Ferrari *et al*. 2019) as well as results from previous studies which used machine learning to identify sequence motifs related to circadian expression in Arabidopsis(Gardiner *et al*. 2021). This provides evidence for the utility of transferring knowledge from fundamental biology findings in model species to inform understanding about molecular biology in crop plants, despite >100M years of evolutionary divergence.

### Co-segregation of promoter haplotypes and transcript rhythmicity

Following the identification of the predictive motifs from our CNN, we hypothesized that sequence variation could be causing differing patterns of diel gene expression among alleles of a gene. For each pangene, we tested for an association between the haplotype sequence of each inbred’s promoter region and patterns of diel transcript abundance, identifying 489 pangenes with significant association between promoter haplotype and diel expression pattern (Figure 2). We refer to these genes from here onward as Differential Diel Regulation (DDR) candidates (Supplemental Files 2 and 3).

**Figure 2:**
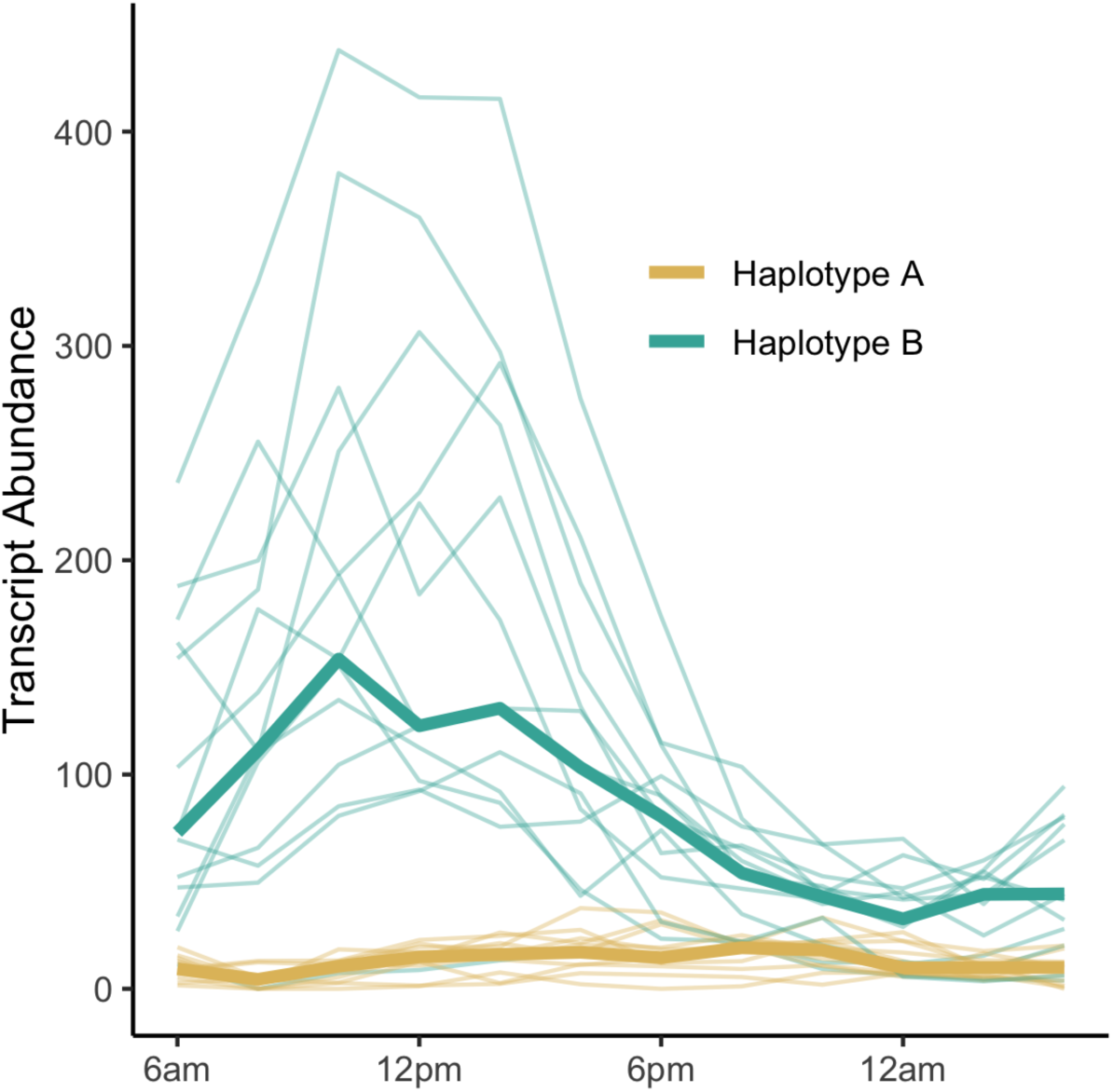
Example of a pangene (Zm00001eb017170; ortholog of Arabidopsis ROQH1) with differential diel regulation based on promoter haplotype. Thin lines show transcript abundance for a single inbred throughout the course of the day and are colored based on which haplotype the inbred has in the promoter region of the gene; thick lines show the median transcript abundance for individuals with each of two haplotypes.

### Function of DDR genes

Among the identified DDR genes are calmodulin2 (cal2; Zm00001eb128040), a calcium signaling gene previously reported as involved in flowering time and adaptation to higher altitudes(Li *et al*. 2016; Hu *et al*. 2022) and MADS-transcription factor 68 (mads68; Zm00001eb006480), an ortholog of rice mads47 implicated in floral development, brassinosteroid signaling, and in maize, endosperm development(Duan *et al*. 2006; Fornara *et al*. 2008; Qiao *et al*. 2016).

KEGG pathway analysis of DDR genes reveals that they are enriched for a number of KEGG pathways related to metabolism (Figure 3). The highest enrichment observed was for the sulfur metabolism pathway, which, along with cysteine and methionine metabolism (fourth highest enrichment), has been previously reported as downregulated in arabidopsis mutants lacking dawn clock components (LHY and CCA1)(Flis *et al*. 2019). We also found enrichment for carbon metabolism and amino acid biosynthesis, which have been shown to be interlinked in diel regulatory loops between photosynthesis, carbon metabolism, and nitrogen metabolism(Stitt *et al*. 2002). Finally, enrichment for ribosomes and protein processing agree with previous work demonstrating that translation of ribosomal proteins is diurnally regulated(Missra *et al*. 2015) and that translation and protein degradation rates are tightly diurnally coupled(Mehta *et al*. 2021; Duncan and Millar 2022). Because plant growth relies on effectively coordinated synthesis and degradation of starch during the day and night(Stitt and Zeeman 2012), respectively, it is quite possible that allelic variation for diel transcription of genes in these various metabolic pathways result from, or contribute to, variable (mal)adaptation among these 24 inbred lines to the diurnal conditions under which they were grown.

**Figure 3:**
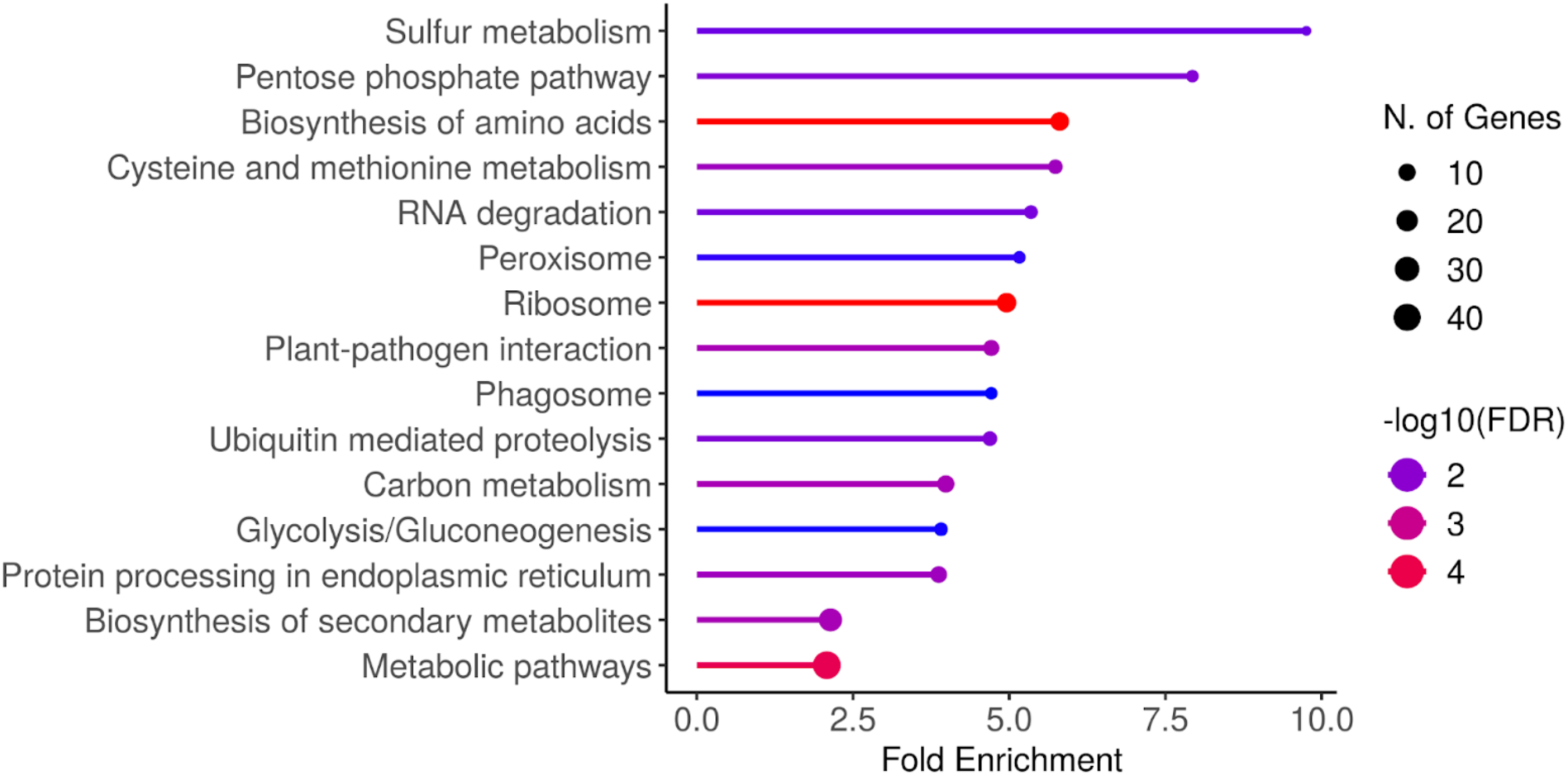
KEGG pathway enrichment results from 489 DDR genes, compared to the background set of 33,201 genes transcribed in at least one tissue of at least one NAM parent line.

### Comparison of diel transcript abundance in an independent experiment

We compared diel patterns of transcript abundance in our field grown plants to three inbreds sampled every three hours in a growth chamber (Supplemental Data 4). We found that among those three inbreds, rank order for the number of significantly cycling genes was preserved. Of all genes that we observed as significantly cycling in the field, 38% (CML103) to 68% (P39) were also significantly cycling in the growth chamber assay (Supplemental Figure 3, Supplemental File 4). This underscores the fact that diel regulation of transcription is consistent across development and environment for some genes, but condition-specific for others. Nearly half of the DDR genes identified in the field experiment were also identified as DDR in the growth chamber, despite the fact that not all haplotypes from the field experiment are represented by the limited genetic diversity of only three inbreds in the growth chamber experiment, as well as the fact that we used a different statistical method for assessing the significance of DDR in growth chamber materials (see Methods). For the following enrichment analyses, we used 205 genes identified as DDR in both field and growth chamber experiments (Supplemental Files 5 and 6).

### DDR candidates are enriched for GWAS hits

Because the circadian clock regulates numerous biological processes(Lu *et al*. 2017), we hypothesized that allelic differences in diel transcription patterns contribute to natural variation for traits regulated by the circadian clock. To test this hypothesis, we checked whether DDR candidates are enriched for GWAS hits relative to randomly selected genes. However, choosing an appropriately matched set of genes from which to take random samples is non-trivial, in order to ensure that the random samples are truly representative and a fair comparison to the DDR genes under consideration.

We compared GWAS hits in DDR genes to random selections from four different subsets of annotated genes: 1) all annotated genes, as a baseline. This null set was likely to overestimate the significance of DDR genes, because the set of all gene annotations contains pseudogenes and incorrectly annotated genes that are likely to have fewer significant GWAS associations. 2) all genes that show evidence of transcription in at least one tissue, in at least one of the 26 NAM parents(Hufford *et al*. 2021). 3) genes that show evidence of diel rhythmicity in at least two inbreds. This gene set was chosen to represent the set of genes that are either segregating for rhythmicity or constitutively rhythmic. 4) genes that show evidence of diel rhythmicity in at least 22 inbreds. This gene set was chosen to represent genes that are constitutively, or nearly constitutively, rhythmic.

We found that DDR genes have significantly enriched GWAS associations for photoperiod sensitivity traits compared to all genes (p<0.001), genes transcribed in at least one tissue/inbred (p<0.001), genes segregating for rhythmicity (p<0.02), and constitutively rhythmic genes, (p<0.004). DDR genes show weaker enrichment for flowering time traits, with significant enrichment only compared to all genes or genes transcribed in at least one tissue/inbred (Figure 4A).

**Figure 4:**
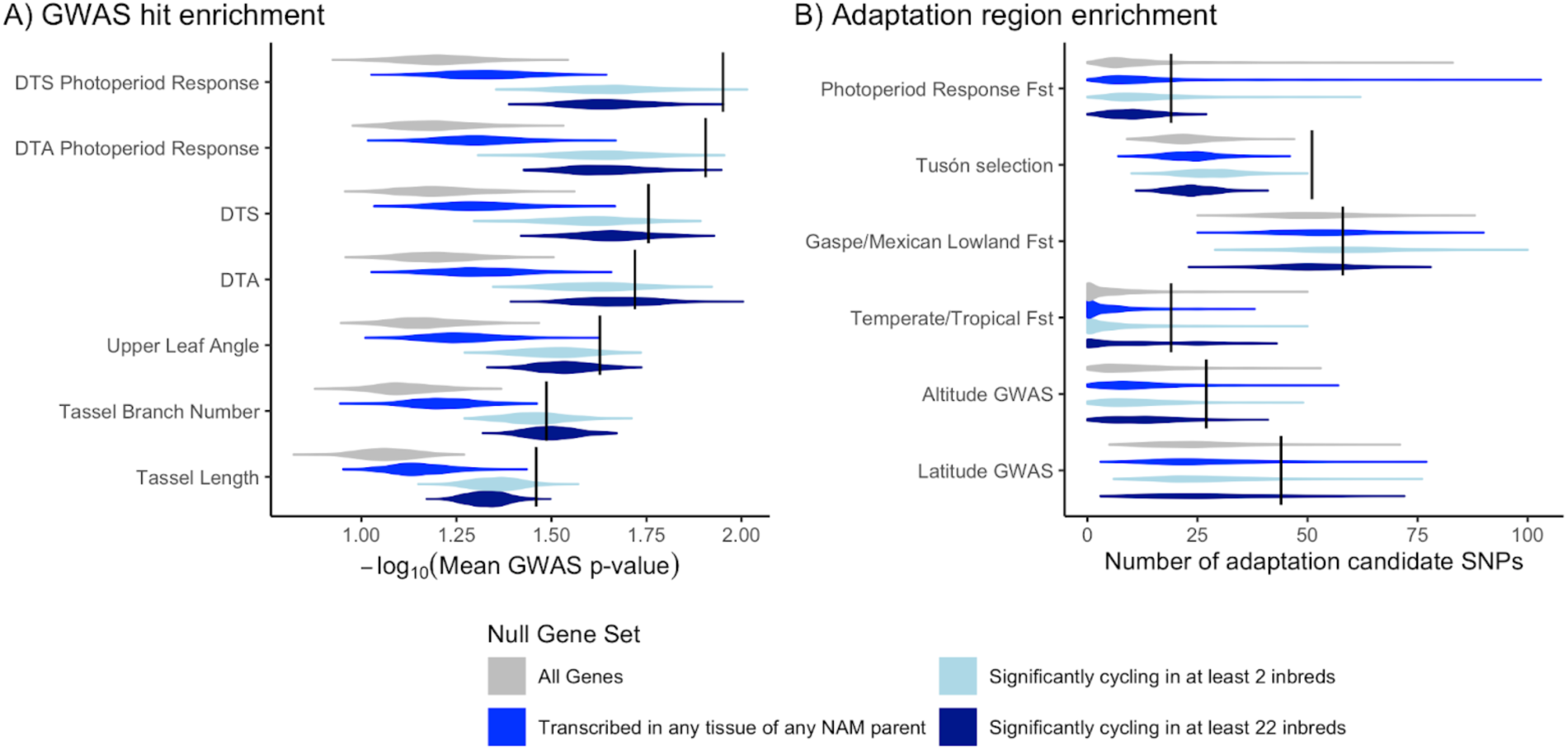
(A) Enrichment for GWAS signal in 205 DDR genes (black vertical lines) compared to random samples of 205 genes from four different null sets (gray and blue distributions). (B) Overlap between candidate adaptation SNPs from five previous scans for selection and DDR genes (black lines). Gray and blue distributions show overlap between candidate adaptation SNPs and randomly selected genes from different null gene sets.

GWAS hits for control traits (tassel length, tassel branch number, and leaf angle; chosen because we predicted they would not be influenced by DDR genes) were also all significantly enriched in DDR genes when compared to all annotated genes (p<0.001) or genes transcribed in at least one tissue/inbred (p<0.002). Compared to genes segregating for rhythmicity and genes with constitutive rhythmicity, DDR genes showed no enrichment in GWAS signal for leaf angle or tassel branch number (p>0.09) (Figure 4A). Overall, the three control traits showed lower mean GWAS p-values at DDR genes than flowering time and photoperiod sensitivity, indicating DDR genes are more strongly associated with photoperiod and flowering than with control traits. We also checked whether DDR genes overlap with previously identified QTL for photoperiod sensitivity(Hung *et al*. 2012b), but found no evidence for a greater rate of overlap than randomly chosen genes (p=0.51).

These findings demonstrate that DDR genes are enriched for GWAS hits related to photoperiod sensitivity, and that GWAS hit enrichment for flowering time, tassel traits, and leaf angle depends on choice of null gene set. Comparing to genes that show evidence of segregating for transcript rhythmicity (null set #2) may be overly conservative: even though many of those genes were not picked up as DDR candidates in our haplotype-based testing, it may that their segregation for rhythmicity contributes to complex trait variation (ie, the null set may be contain false negative DDR genes). On the other hand, comparing to genes that show any evidence of transcription may be an overly liberal test because the null set contains genes that are transcribed but not relevant to variation in the traits that we tested. The fact that DDR candidates show stronger GWAS associations than control genes for flowering time and photoperiod sensitivity traits support the hypothesis that allelic variation for diel transcription patterns contributes to longitudinal adaptation to longer days.

### Night-Day eQTL are enriched for GWAS hits

Next, using an independent dataset we tested our hypothesis that variable cis-regulation of diel transcription patterns contributes to natural variation for traits regulated by the circadian clock. Using RNAseq data from mature leaf tissue sampled in the Goodman Association Panel (GAP)(Kremling *et al*. 2018), we calculated the difference between nighttime and daytime transcript abundances as a rough estimator for how much transcript abundance varies diurnally, and used the difference as the response variable for eQTL mapping.

We found that Night-Day cis-eQTL are highly enriched in GWAS hits for flowering time and photoperiod response for days to anthesis, compared to random SNPs sampled from within the same genomic context (<5,000bp away from the focal gene) and matched for minor allele frequency (Figure 5). There was no enrichment for tassel traits, as expected, or days to silk photoperiod response, which was unexpected given the strong enrichment for days to anthesis photoperiod GWAS hits. These results also support the idea that cis-regulation of diel transcription may be contributing to changes in the adaptive traits of flowering time and photoperiod sensitivity.

**Figure 5:**
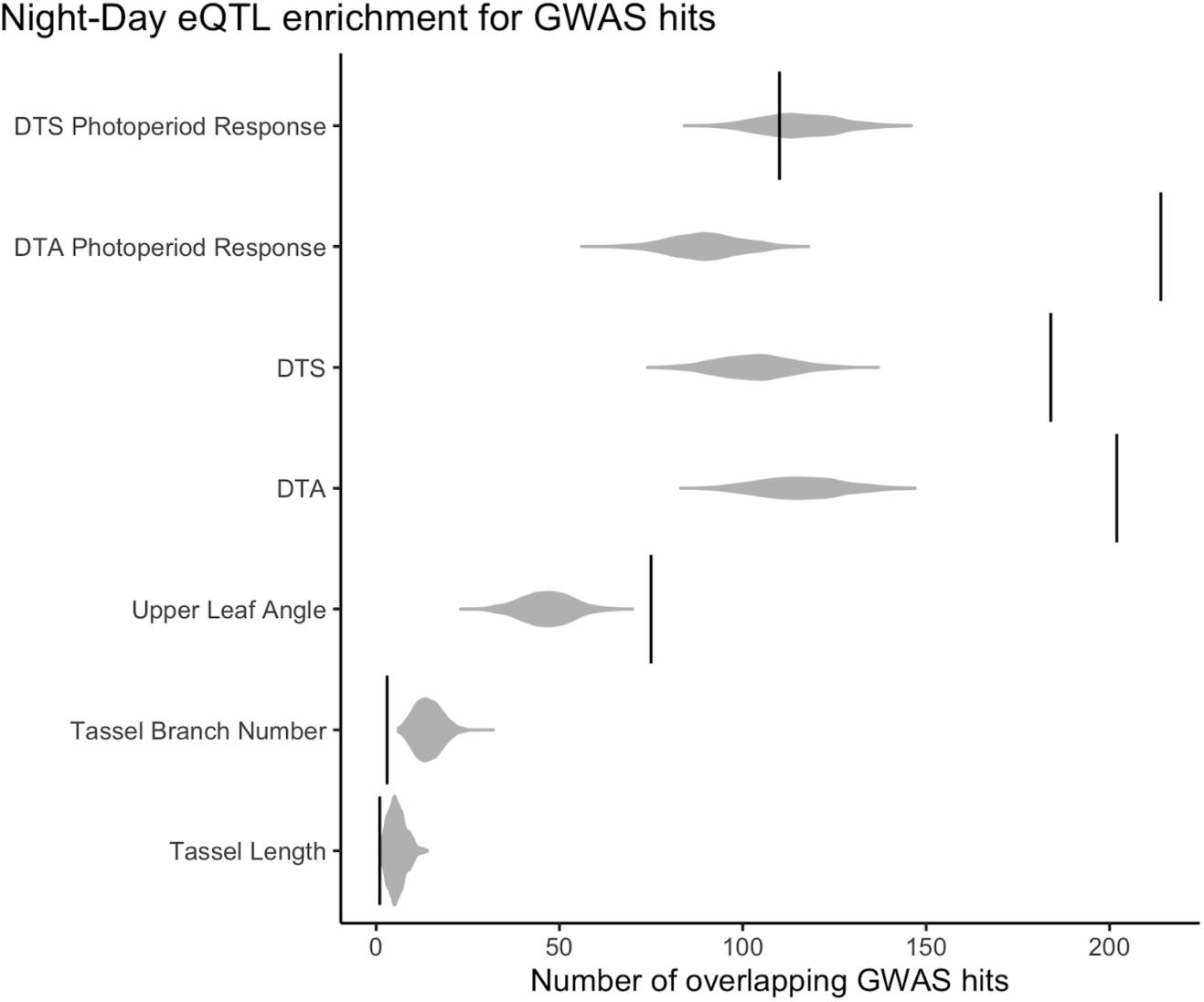
Overlap between GWAS hits and cis-eQTL for the difference between daytime and nighttime transcript abundance (black vertical lines). Gray distributions represent overlap between GWAS hits and SNPs randomly sampled while controlling for minor allele frequency and distance from the nearest gene.

### DDR candidates are moderately enriched for signatures of selection

Given that DDR candidates are enriched for GWAS hits related to flowering time and photoperiod sensitivity, and that cis-regulatory sequences are likely to evolve faster than TFs, changing gene regulation and ultimately phenotype(Stone and Wray 2001; Weirauch and Hughes 2010), we hypothesized that diel transcription patterns may have been subjected to selection during adaptation of maize to higher latitudes. Using several independent studies of maize adaptation to changes in latitude and altitude(Romero Navarro *et al*. 2017; Swarts *et al*. 2017; Gage *et al*. 2017; Wisser *et al*. 2019), we tested DDR candidates for enrichment of putatively selected regions (Figure 4B).

Unlike our GWAS enrichment results, tests for enrichment of adaptive regions among DDR genes are not as sensitive to choice of null set; results remain similar regardless of whether we selected random samples from all genes, genes that show any evidence of transcription, or genes that show evidence of rhythmic transcription.

DDR candidates are significantly enriched (p<0.001) for SNPs showing evidence of selection in Hallauer’s Tusón, a population of tropical traditional varieties that were selected for early flowering in Iowa, USA for 10 generations(Teixeira *et al*. 2015; Hallauer and Carena 2016). We observed low to no significance, however, for enrichment of putative adaptation SNPs identified from Fst between high and low photoperiod sensitivity lines or from three other published studies: eGWAS for latitude or altitude of origin(Romero Navarro *et al*. 2017) (p-value 0.04-0.12), Fst between mexican lowland and northern flint accessions(Swarts *et al*. 2017) (p-value 0.24-0.51), and Fst between 30 tropical and 30 temperate inbreds(Gage *et al*. 2017) (p-value 0.03-0.22).

Adaptation is complex, and involves modulation of processes beyond just those related to flowering and photoperiod sensitivity. It is possible that we were able to observe an enriched overlap of DDR genes and selected loci in Hallauer’s Tusón because the population was directly selected for early flowering time in a single location for ten generations, a less complicated selection history than experienced by natural populations. In the two studies of Fst between temperate and tropical accessions, the lack of enrichment for signals of selection near DDR genes may be because Fst is detecting allele frequency changes related to traits other than photoperiod sensitivity or flowering, or false positives in alleles that have had many more than ten generations to drift. The Romero Navarro study of latitude and altitude was limited in its representation of traditional varieties from latitudes that would induce a photoperiod response(Romero Navarro *et al*. 2017).

Although there is limited statistical significance for these tests of enrichment, in all instances the DDR genes had greater overlap with selection candidates than the median of all permutations, and across all tests the mean enrichment of selection candidates was 275% over the median value from permutations. P-values from Fisher’s combined test for p-values across all selection experiments ranged from 1.4e-5 (all genes) to 9.5e-5 (significantly cycling in two or more inbreds). These results support the hypothesis that DDR genes have been the subject of selection during adaptation to new environments, though the strength and consistency selection seems variable depending on the population.

## Conclusions

In this study, we show that allelic variation for diel patterns of transcript abundance are common in maize and differences in diel transcript abundance rhythms appear at least partially explainable by sequence variation in cis-regulatory regions. It is also possible that transcriptional differences observed in this study are caused by differential chromatin accessibility or variation for trans-acting mechanisms. Observed variation in diel patterns of transcript abundance appear to influence photoperiod sensitivity (phenotypic plasticity for flowering time) and may have been targets of selection when maize was adapted to longer day lengths. These results are an example of how selection on sequence variation impacting gene expression can contribute to phenotypic plasticity and adaptation to new environments.

## Materials and Methods

### Germplasm and field experiment

We chose 24 of the 26 parents of the maize nested association panel (NAM)(Yu *et al*. 2008; Gage *et al*. 2020) to assay for diel patterns of transcript abundance. Inbreds were grown in the field at the Musgrave Research Farm in Aurora, NY in the summer of 2019. Due to different rates of development between inbreds, we sampled tissue from the first leaf without epicuticular wax to standardize the developmental stage of sampled tissue (Supplemental File 7). The last leaf with epicuticular wax was scored on July 16, 2019. Tissue was collected every two hours from 6am on July 19, 2019 until 4am on July 20, 2019. Approximately 1 cm^2^ of leaf tissue was collected from the center of the leaf blade, roughly measured by bending the tip back to the leaf base, on one side of the midrib. To avoid assaying wounding response, a different plant was sampled at each timepoint. If not enough plants were present to sample a different plant at each timepoint, a small number of timepoints were sampled from previously sampled plants, but from the opposite side of the midrib. Tissue samples were collected onto liquid nitrogen and kept cold throughout the experiment on dry ice.

### Growth chamber experiment

Four genotypes (three of which overlap with genotypes grown in the field), B73, Mo17, P39, and CML103, were sown in 24-cell trays in 1:1 Lambert GPM:Turface. Two pots of each genotype were planted for each of 11 timepoints (0h through 30h), except for the 12h timepoint, for which four pots of each genotype were planted. Each pot was planted with 2 seeds on April 9, 2019. An additional 44 pots were planted in the same manner with B73 to be used as a border in the growth chamber. Experimental (non-border) pots were randomized every 2-3 days to reduce spatial effects. The growth chamber was set to 14-hour days at 26°C and 10 hour nights at 22°C.

Tissue sampling occurred when plants were at approximately v3 growth stage, younger than plants sampled in the field experiment. Tissue was collected from the middle of the second leaf from the healthiest plant in each pot starting on April 22, 2019 at 09:00 (∼1 hour after lights come on) and continuing every 3 hours until April 23, 2019 at 15:00. Tissue samples were collected onto liquid nitrogen and kept cold throughout the experiment on dry ice.

### RNA-seq and bioinformatic analysis

RNA extraction and 3’ RNA sequencing was performed by the Genomics Facility at the Cornell Institute of Biotechnology following methods previously described(Kremling *et al*. 2018). Cutadapt(Martin 2011) version 2.3 was used to trim 12 bases from the 3’ end as recommended by Lexogen, remove bases with quality <20 from the 3’ end, remove TruSeq adapters and any poly-A or poly-Ts. Because all the NAM parents have de novo genome assemblies(Hufford *et al*. 2021), read alignment and transcript abundance estimation was performed using the cognate genome assembly for each of the 24 inbred lines to reduce reference bias. Read alignment and counting wer done using Salmon(Patro *et al*. 2017) v1.8 with the options --libType A --validateMappings --noLengthCorrection. Transcript abundance was normalized using the estimateSizeFactors() and counts() functions from the R(R Core Team 2018) package DESeq2(Love *et al*. 2014).

### Predicting cycling status from promoter sequence

For the field experiment data, each gene in each inbred was evaluated for rhythmicity in R(R Core Team 2018) using the MetaCycle(Wu *et al*. 2016) function *meta2d()*, as well as the diffCircadian(Ding *et al*. 2021) function *LR_rhythmicity()*. Because the *meta2d()* function aggregates rhythmicity test statistics from three different methods, using both MetaCycle and diffCircadian resulted in testing each gene for rhythmic patterns of transcript abundance via four different methods.

To model the relationship between promoter sequence and rhythmic diel patterns of transcript abundance, we adapted a modification of the DanQ model(Quang and Xie 2016), originally developed to predict chromatin accessibility. The model architecture consists of convolutional and max-pooling layers followed by a bidirectional LSTM to allow the model to learn DNA sequence motifs and their interactions that are predictive of cyclical transcript abundance patterns.

Our model was trained with the sequence from 1000bp upstream to 500bp downstream of the annotated transcription start site of the first transcript (“T001”) of each gene as features. Each gene had three continuous labels: relative amplitude computed by MetaCycle, and p-value and R^2^ from diffCircadian. We chose these labels to attempt to quantitatively capture the amount of evidence for rhythmic diel transcript abundance patterns.

We trained our model for 500 epochs, using all genes from 12 inbreds as training data and all genes from 3 inbreds as validation data. We converted the trained convolutional filters into sequence motifs using methods adapted from(Koo and Eddy 2019). Filter sequence logos were inspected visually and compared to known, previously published Arabidopsis TF binding motifs.

### Identification of DDR candidates in field experiment

Using the maize PHG v1 (available at https://phg.maizegdb.org, accessed September 2024)(Bradbury *et al*. 2022) method name “CONSENSUS_84plusRef_mxDiv_10toNeg3_maxClusers30”, we assigned haplotypes for reference ranges that overlapped with the 1000bp upstream of each gene in each inbred. For each pangene, we tested for an association between haplotype ID and the difference in magnitude between maximum and minimum transcript abundance during the 24 hour time course using a Kruskal-Wallis (KW) test. To estimate p-values, we permuted haplotype labels and re-calculated the KW test statistic 10,000 times per reference range.

Pangenes were considered to be Differential Diel Regulation (DDR) candidates if the p-value of the KW permutation test was < 0.05, at least six inbreds had significant single-gene cycling tests (p < 0.01), and at least six inbreds had baseline transcript abundance values greater than 50. The additional filtering on single-gene cycling tests ensured that the KW test of change in transcript abundance was including genes with some evidence of diel rhythmicity, and the baseline transcript abundance filtering was to remove noisy, low-expressed pangenes.

### KEGG pathway enrichment

Gene ontology enrichment was performed using ShinyGO v0.8.1(Ge *et al*. 2020) (https://bioinformatics.sdstate.edu/go/, accessed December 6, 2024). The web application was used with the Ensembl/STRING-db ID set to “zmays_eg_gene”. The 489 DDR genes were used as input, and the background was set to the list of all genes transcribed in at least one NAM parent, in at least one tissue (33,201 genes). All other settings were left at default, and we evaluated results for KEGG pathways.

### Identification of DDR genes in growth chamber experiment

To robustly filter the DDR genes identified in the field experiment, we tested the 489 field experiment DDR genes for allelic differences in diel transcription in the growth chamber experiment. We grouped P39, B73, and CML103 samples (each genotype replicated twice per timepoint) based on haplotype imputed from the PHG, as described for the field experiment data above. We then tested for differences between haplotypes using the *LR_diff()* function from the diffCircadian package(Ding *et al*. 2021). If all three genotypes had the same promoter haplotype, the gene was not tested. If all three genotypes belonged to two haplotype groups, we simply tested the difference between the two haplotypes. If all three genotypes had three different haplotypes, we performed pairwise tests between all three haplotypes and kept the lowest p-value. 205 genes with a p-value for differences in amplitude less than 0.01 were considered significant and used for GWAS and adaptation enrichment experiments below.

### Enrichment for GWAS results and evidence of adaptation

We performed GWAS in the maize Goodman Association Panel (GAP) population, which consists of 282 diverse inbred maize lines(Flint-Garcia *et al*. 2005). Phenotypic data were previously collected (Tian *et al*. 2011; Hung *et al*. 2012a; b) and compiled for ease of use(Khaipho-Burch *et al*. 2023). We followed the methods for fast association described in(Khaipho-Burch *et al*. 2023), using high density genotypes for the GAP on the B73v5 reference genome (Grzybowski *et al*. 2023). We chose to test for enrichment of GWAS signal in traits related to flowering time and latitudinal adaptation: growing degree days to flowering for silk (DTS) and anthesis (DTA), and photoperiod response, or the difference between DTS (or DTA) under long days and short days. We also chose to include leaf angle, tassel length, and tassel branch number as control traits that we predicted would not be enriched for GWAS signal in DDR candidates. GWAS was performed using rTASSEL(Monier *et al*. 2022), an R-based front end for TASSEL(Bradbury *et al*. 2007).

To compare GWAS p-values from 205 DDR candidates to random samples, we first took the mean of the minimum p-value within each DDR candidate gene +/-1000bp. This was compared to 1,000 random samples of 205 of genes drawn from a null set. Because results of permutation tests like this can be highly dependent on the choice of null set, we tested four different null sets: 1) all annotated gene models; 2) annotated genes with >5 transcripts per million (TPM) in at least one tissue of at least one NAM parent(Hufford *et al*. 2021); 3) annotated genes that show evidence of cycling (p<0.01) in at least two inbreds in this study; 4) annotated genes that show evidence of cycling (p<0.01) in 22 or more inbreds in this study. The second null set was chosen to reduce our sampling of pseudogenes or other non-functional genes relative to the first null set. Null sets #3 and #4 were chosen to capture genes showing segregation for rhythmicity and near-constitutive rhythmicity, respectively.

Next, we tested whether DDR candidates showed evidence of enrichment for signatures of selection or adaptive function. SNP variants showing evidence of selection or adaptation were collected from previous studies of: environmental GWAS of maize landrace adaptation to latitude and altitude(Romero Navarro *et al*. 2017); F_ST_ between temperate and tropical maize inbreds(Gage *et al*. 2017); F_ST_ between maize from central Mexican lowlands and Gaspé Flint from Quebec(Swarts *et al*. 2017); and directed, short term evolution in the maize Tuson population for day length adaptation(Wisser *et al*. 2019). In addition, we calculated Fst between 30 inbred lines with the lowest photoperiod response values and 30 inbred lines with the highest photoperiod response values in the GAP(Hung *et al*. 2012b) using SNP data from (Grzybowski *et al*. 2023) in VCFtools(Danecek *et al*. 2011) and kept the top 10,000 SNPs as putative adaptation variants.

For each set of adaptation candidate SNPs, we first counted the number of adaptation SNPs that overlapped with DDR genes +/-1000bp. We then sampled an equal number of genes from each of our four null sets and counted the number of adaptation candidate SNPs that overlapped with this sample. This sampling procedure was repeated 1,000 times to generate a null distribution, and the p-value for each set of DDR candidates was estimated as the proportion of random samples that had more overlapping adaptation SNPs than the DDR candidate genes.

### eQTL overlap with GWAS hits

We performed an additional test for whether changes in cis-regulatory sequences associated with changes in transcript abundance co-localize with GWAS hits for photoperiod sensitivity, flowering time, and control traits. Using daytime and nighttime leaf transcript abundance data from(Kremling *et al*. 2018), we estimated the degree of diel fluctuation in transcripts by subtracting estimates of daytime expression from estimates of nighttime expression. We then performed eQTL mapping in 203 individuals using the same techniques described by(Kremling *et al*. 2018). We then counted the overlaps between cis-eQTL, within 5,000bp of their focal gene, and GWAS hits (variants with p-value < 1e-5) for photoperiod sensitivity, flowering time, leaf angle, tassel length, and tassel branch number. This overlap was compared to 10,000 random draws of the same number of SNPs as cis-eQTL, matched for distance to the nearest gene and distribution of minor allele frequency.

## Code availability

Code will be available upon final publication at https://github.com/joegage/diverse_diel

## Data availability

All sequencing data will be available on NCBI SRI upon final publication. All supplementary data will be made publicly available upon final publication.

## Acknowledgements

This research was supported in part by the intramural research program of the U.S. Department of Agriculture, National Institute of Food and Agriculture Hatch 7002327. Research reported in this publication was supported by the USDA-ARS, National Institute of General Medical Sciences of the National Institutes of Health under award number R35GM151048, and NSF Grant IOS-1906619. Sequencing was performed by the Biotechnology Resource Center (BRC) Genomics Facility (RRID:SCR_021727) at the Cornell Institute of Biotechnology. Computing was supported by Cornell University BRC Bioinformatics Core Facility (RRID:SCR_021757) and North Carolina State University High Performance Computing Services Core Facility (RRID:SCR_022168).

## References

Bradbury P. J., Z. Zhang, D. E. Kroon, T. M. Casstevens, Y. Ramdoss, et al., 2007 TASSEL: software for association mapping of complex traits in diverse samples. Bioinformatics 23: 2633–2635.

Bradbury P. J., T. Casstevens, S. E. Jensen, L. C. Johnson, Z. R. Miller, et al., 2022 The Practical Haplotype Graph, a platform for storing and using pangenomes for imputation. Bioinformatics 38: 3698–3702.

Buckler E. S., J. B. Holland, P. J. Bradbury, C. B. Acharya, P. J. Brown, et al., 2009 The genetic architecture of maize flowering time. Science 325: 714–718.

Crawford G. W., D. Saunders, and D. G. Smith, 2006 Pre-contact maize from Ontario, Canada: Context, chronology, variation, and plant association. Histories of maize: Multidisciplinary approaches to the prehistory, linguistics, biogeography, domestication, and evolution of maize 549–559.

Danecek P., A. Auton, G. Abecasis, C. A. Albers, E. Banks, et al., 2011 The variant call format and VCFtools. Bioinformatics 27: 2156–2158.

Ding H., L. Meng, A. C. Liu, M. L. Gumz, A. J. Bryant, et al., 2021 Likelihood-based tests for detecting circadian rhythmicity and differential circadian patterns in transcriptomic applications. Brief. Bioinform. 22. 10.1093/bib/bbab224

Duan K., L. Li, P. Hu, S.-P. Xu, Z.-H. Xu, et al., 2006 A brassinolide-suppressed rice MADS-box transcription factor, OsMDP1, has a negative regulatory role in BR signaling. Plant J. 47: 519–531.

Duncan O., and A. H. Millar, 2022 Day and night isotope labelling reveal metabolic pathway specific regulation of protein synthesis rates in Arabidopsis. Plant J. 109: 745–763.

Ferrari C., S. Proost, M. Janowski, J. Becker, Z. Nikoloski, et al., 2019 Kingdom-wide comparison reveals the evolution of diurnal gene expression in Archaeplastida. Nat. Commun. 10: 737.

Flint-Garcia S. A., A.-C. Thuillet, J. Yu, G. Pressoir, S. M. Romero, et al., 2005 Maize association population: a high-resolution platform for quantitative trait locus dissection: High-resolution maize association population. Plant J. 44: 1054–1064.

Flis A., V. Mengin, A. A. Ivakov, S. T. Mugford, H.-M. Hubberten, et al., 2019 Multiple circadian clock outputs regulate diel turnover of carbon and nitrogen reserves: Metabolism in five circadian clock mutants. Plant Cell Environ. 42: 549–573.

Fornara F., V. Gregis, N. Pelucchi, L. Colombo, and M. Kater, 2008 The rice StMADS11-like genes OsMADS22 and OsMADS47 cause floral reversions in Arabidopsis without complementing the svp and agl24 mutants. Journal of Experimental Botany 59: 2181– 2190.

Gage J. L., D. Jarquin, C. Romay, A. Lorenz, E. S. Buckler, et al., 2017 The effect of artificial selection on phenotypic plasticity in maize. Nat. Commun. 8: 1348.

Gage J. L., B. Monier, A. Giri, and E. S. Buckler, 2020 Ten Years of the maize Nested Association Mapping Population: Impact, Limitations, and Future Directions. Plant Cell. 10.1105/tpc.19.00951

Gardiner L.-J., R. Rusholme-Pilcher, J. Colmer, H. Rees, J. M. Crescente, et al., 2021 Interpreting machine learning models to investigate circadian regulation and facilitate exploration of clock function. Proc. Natl. Acad. Sci. U. S. A. 118: e2103070118.

Ge S. X., D. Jung, and R. Yao, 2020 ShinyGO: a graphical gene-set enrichment tool for animals and plants. Bioinformatics 36: 2628–2629.

Gendron J. M., J. L. Pruneda-Paz, C. J. Doherty, A. M. Gross, S. E. Kang, et al., 2012 *Arabidopsis* circadian clock protein, TOC1, is a DNA-binding transcription factor. Proc. Natl. Acad. Sci. U. S. A. 109: 3167–3172.

Gnesutta N., R. W. Kumimoto, S. Swain, M. Chiara, C. Siriwardana, et al., 2017 CONSTANS Imparts DNA Sequence Specificity to the Histone Fold NF-YB/NF-YC Dimer. Plant Cell 29: 1516–1532.

Grzybowski M. W., R. V. Mural, G. Xu, J. Turkus, J. Yang, et al., 2023 A common resequencing-based genetic marker data set for global maize diversity. Plant J. 113: 1109–1121.

Hallauer A. R., and M. J. Carena, 2016 Registration of BS39 maize germplasm. J. Plant Regist. 10: 296–300.

Hu H., T. Crow, S. Nojoomi, A. J. Schulz, J. M. Estévez-Palmas, et al., 2022 Allele-specific expression reveals multiple paths to highland adaptation in maize. Mol. Biol. Evol. 39: msac239.

Hufford M. B., A. S. Seetharam, M. R. Woodhouse, K. M. Chougule, S. Ou, et al., 2021 De novo assembly, annotation, and comparative analysis of 26 diverse maize genomes. Science 373: 655–662.

Hung H.-Y., C. Browne, K. Guill, N. Coles, M. Eller, et al., 2012a The relationship between parental genetic or phenotypic divergence and progeny variation in the maize nested association mapping population. Heredity 108: 490–499.

Hung H.-Y., L. M. Shannon, F. Tian, P. J. Bradbury, C. Chen, et al., 2012b ZmCCT and the genetic basis of day-length adaptation underlying the postdomestication spread of maize. Proc. Natl. Acad. Sci. U. S. A. 109: E1913–21.

Khaipho-Burch M., T. Ferebee, A. Giri, G. Ramstein, B. Monier, et al., 2023 Elucidating the patterns of pleiotropy and its biological relevance in maize. PLoS Genet. 19: e1010664.

Khan S., S. C. Rowe, and F. G. Harmon, 2010 Coordination of the maize transcriptome by a conserved circadian clock. BMC Plant Biol. 10: 126.

Koo P. K., and S. R. Eddy, 2019 Representation learning of genomic sequence motifs with convolutional neural networks. PLoS Comput. Biol. 15: e1007560.

Kremling K. A. G., S.-Y. Chen, M.-H. Su, N. K. Lepak, M. C. Romay, et al., 2018 Dysregulation of expression correlates with rare-allele burden and fitness loss in maize. Nature 555: 520–523.

Lambert S. A., A. Jolma, L. F. Campitelli, P. K. Das, Y. Yin, et al., 2018 The Human Transcription Factors. Cell 172: 650–665.

Li Y.-X., C. Li, P. J. Bradbury, X. Liu, F. Lu, et al., 2016 Identification of genetic variants associated with maize flowering time using an extremely large multi-genetic background population. Plant J. 86: 391–402.

Love M. I., W. Huber, and S. Anders, 2014 Moderated estimation of fold change and dispersion for RNA-seq data with DESeq2. Genome Biol. 15: 550.

Lu H., C. R. McClung, and C. Zhang, 2017 Tick Tock: Circadian Regulation of Plant Innate Immunity. Annu. Rev. Phytopathol. 55: 287–311.

Martin M., 2011 Cutadapt removes adapter sequences from high-throughput sequencing reads. EMBnet.journal 17: 10–12.

Matsuoka Y., Y. Vigouroux, M. M. Goodman, J. Sanchez G, E. Buckler, et al., 2002 A single domestication for maize shown by multilocus microsatellite genotyping. Proc. Natl. Acad. Sci. U. S. A. 99: 6080–6084.

Mehta D., J. Krahmer, and R. G. Uhrig, 2021 Closing the protein gap in plant chronobiology. Plant J. 106: 1509–1522.

Michael T. P., P. A. Salomé, H. J. Yu, T. R. Spencer, E. L. Sharp, et al., 2003 Enhanced fitness conferred by naturally occurring variation in the circadian clock. Science 302: 1049– 1053.

Michael T. P., 2022 Core circadian clock and light signaling genes brought into genetic linkage across the green lineage. Plant Physiol. 190: 1037–1056.

Missra A., B. Ernest, T. Lohoff, Q. Jia, J. Satterlee, et al., 2015 The circadian clock modulates global daily cycles of mRNA ribosome loading. Plant Cell 27: 2582–2599.

Monier B., T. M. Casstevens, P. J. Bradbury, and E. S. Buckler, 2022 rTASSEL: An R interface to TASSEL for analyzing genomic diversity. J. Open Source Softw. 7: 4530.

Nagel D. H., C. J. Doherty, J. L. Pruneda-Paz, R. J. Schmitz, J. R. Ecker, et al., 2015 Genome-wide identification of CCA1 targets uncovers an expanded clock network in Arabidopsis. Proc. Natl. Acad. Sci. U. S. A. 112: E4802–10.

Patro R., G. Duggal, M. I. Love, R. A. Irizarry, and C. Kingsford, 2017 Salmon provides fast and bias-aware quantification of transcript expression. Nat. Methods 14: 417–419.

Petersen J., A. Rredhi, J. Szyttenholm, and M. Mittag, 2022 Evolution of circadian clocks along the green lineage. Plant Physiol. 190: 924–937.

Piperno D. R., A. J. Ranere, I. Holst, J. Iriarte, and R. Dickau, 2009 Starch grain and phytolith evidence for early ninth millennium B.P. maize from the Central Balsas River Valley, Mexico. Proc. Natl. Acad. Sci. U. S. A. 106: 5019–5024.

Qiao Z., W. Qi, Q. Wang, Y. Feng, Q. Yang, et al., 2016 ZmMADS47 regulates Zein gene transcription through interaction with Opaque2. PLoS Genet. 12: e1005991.

Quang D., and X. Xie, 2016 DanQ: a hybrid convolutional and recurrent deep neural network for quantifying the function of DNA sequences. Nucleic Acids Res. 44: e107.

R Core Team, 2018 R: A Language and Environment for Statistical Computing

Romero Navarro J. A., M. Willcox, J. Burgueño, C. Romay, K. Swarts, et al., 2017 A study of allelic diversity underlying flowering-time adaptation in maize landraces. Nat. Genet. 49: 476–480.

Sanchez S. E., and S. A. Kay, 2016 The Plant Circadian Clock: From a Simple Timekeeper to a Complex Developmental Manager. Cold Spring Harb. Perspect. Biol. 8. 10.1101/cshperspect.a027748

Sauer J. D., and P. Weatherwax, 1955 Indian corn in old America. Geogr. Rev. 45: 446.

Spensley M., J.-Y. Kim, E. Picot, J. Reid, S. Ott, et al., 2009 Evolutionarily conserved regulatory motifs in the promoter of the Arabidopsis clock gene LATE ELONGATED HYPOCOTYL. Plant Cell 21: 2606–2623.

Stewart A. J., S. Hannenhalli, and J. B. Plotkin, 2012 Why transcription factor binding sites are ten nucleotides long. Genetics 192: 973–985.

Stitt M., C. Müller, P. Matt, Y. Gibon, P. Carillo, et al., 2002 Steps towards an integrated view of nitrogen metabolism. J. Exp. Bot. 53: 959–970.

Stitt M., and S. C. Zeeman, 2012 Starch turnover: pathways, regulation and role in growth. Curr. Opin. Plant Biol. 15: 282–292.

Stone J. R., and G. A. Wray, 2001 Rapid evolution of cis-regulatory sequences via local point mutations. Mol. Biol. Evol. 18: 1764–1770.

Swarts K., R. M. Gutaker, B. Benz, M. Blake, R. Bukowski, et al., 2017 Genomic estimation of complex traits reveals ancient maize adaptation to temperate North America. Science 357: 512–515.

Teixeira J. E. C., T. Weldekidan, N. de Leon, S. Flint-Garcia, J. B. Holland, et al., 2015 Hallauer’s Tusón: a decade of selection for tropical-to-temperate phenological adaptation in maize. Heredity (Edinb.) 114: 229–240.

Tian F., P. J. Bradbury, P. J. Brown, H. Hung, Q. Sun, et al., 2011 Genome-wide association study of leaf architecture in the maize nested association mapping population. Nat. Genet. 43: 159–162.

Torii K., K. Inoue, K. Bekki, K. Haraguchi, M. Kubo, et al., 2022 A guiding role of the Arabidopsis circadian clock in cell differentiation revealed by time-series single-cell RNA sequencing. Cell Rep. 40: 111059.

Troyer A. F., and L. G. Hendrickson, 2007 Background and importance of ‘Minnesota 13’ corn. Crop Sci. 47: 905–914.

Tu X., M. K. Mejía-Guerra, J. A. Valdes Franco, D. Tzeng, P.-Y. Chu, et al., 2020 Reconstructing the maize leaf regulatory network using ChIP-seq data of 104 transcription factors. Nat. Commun. 11: 5089.

Vandepoele K., K. Vlieghe, K. Florquin, L. Hennig, G. T. S. Beemster, et al., 2005 Genome-wide identification of potential plant E2F target genes. Plant Physiol. 139: 316–328.

Wang P., L. Wang, L. Zhang, T. Wu, B. Sun, et al., 2022 Genomic Dissection and Diurnal Expression Analysis Reveal the Essential Roles of the PRR Gene Family in Geographical Adaptation of Soybean. Int. J. Mol. Sci. 23. 10.3390/ijms23179970

Weirauch M. T., and T. R. Hughes, 2010 Conserved expression without conserved regulatory sequence: the more things change, the more they stay the same. Trends Genet. 26: 66–74.

Wisser R. J., Z. Fang, J. B. Holland, J. E. C. Teixeira, J. Dougherty, et al., 2019 The Genomic Basis for Short-Term Evolution of Environmental Adaptation in Maize. Genetics 213: 1479–1494.

Wu G., R. C. Anafi, M. E. Hughes, K. Kornacker, and J. B. Hogenesch, 2016 MetaCycle: an integrated R package to evaluate periodicity in large scale data. Bioinformatics 32: 3351– 3353.

Yu J., J. B. Holland, M. D. McMullen, and E. S. Buckler, 2008 Genetic design and statistical power of nested association mapping in maize. Genetics 178: 539–551.

