## Supplement for "Maize inbreds show allelic variation for diel transcription patterns"

**Supplemental Files and Data**

- Supplemental File 1: Single gene rhythmicity test results from field study
- Supplemental File 2: Haplotype-based test for differences in transcript abundance from field study data
- Supplemental File 3: List of DDR pan-genes identified from field study
- Supplemental File 4: Signe gene rhythmicity test results from growth chamber study
- Supplemental File 5: Haplotype-based test for differences in transcript abundance from growth chamber data
- Supplemental File 6: List of DDR pan-genes identified from field study and growth chamber study
- Supplemental File 7: Details on last leaf with epicuticular wax, and leaf stage at the time of sampling for each inbred in the field study
- Supplemental Data 1: Transcript abundance values for field study
- Supplemental Data 2: Trained model weights for predicting rhythmicity from promoter sequence
- Supplemental Data 3: Sequence motifs learned by the model for predicting rhythmicity from promoter sequence
- Supplemental Data 4: Transcript abundance values for growth chamber study

**Supplemental Figures**

*Supplemental Figure 1: regressions of number cycling genes against flowering time, developmental stage, photoperiod sensitivity, and read depth*

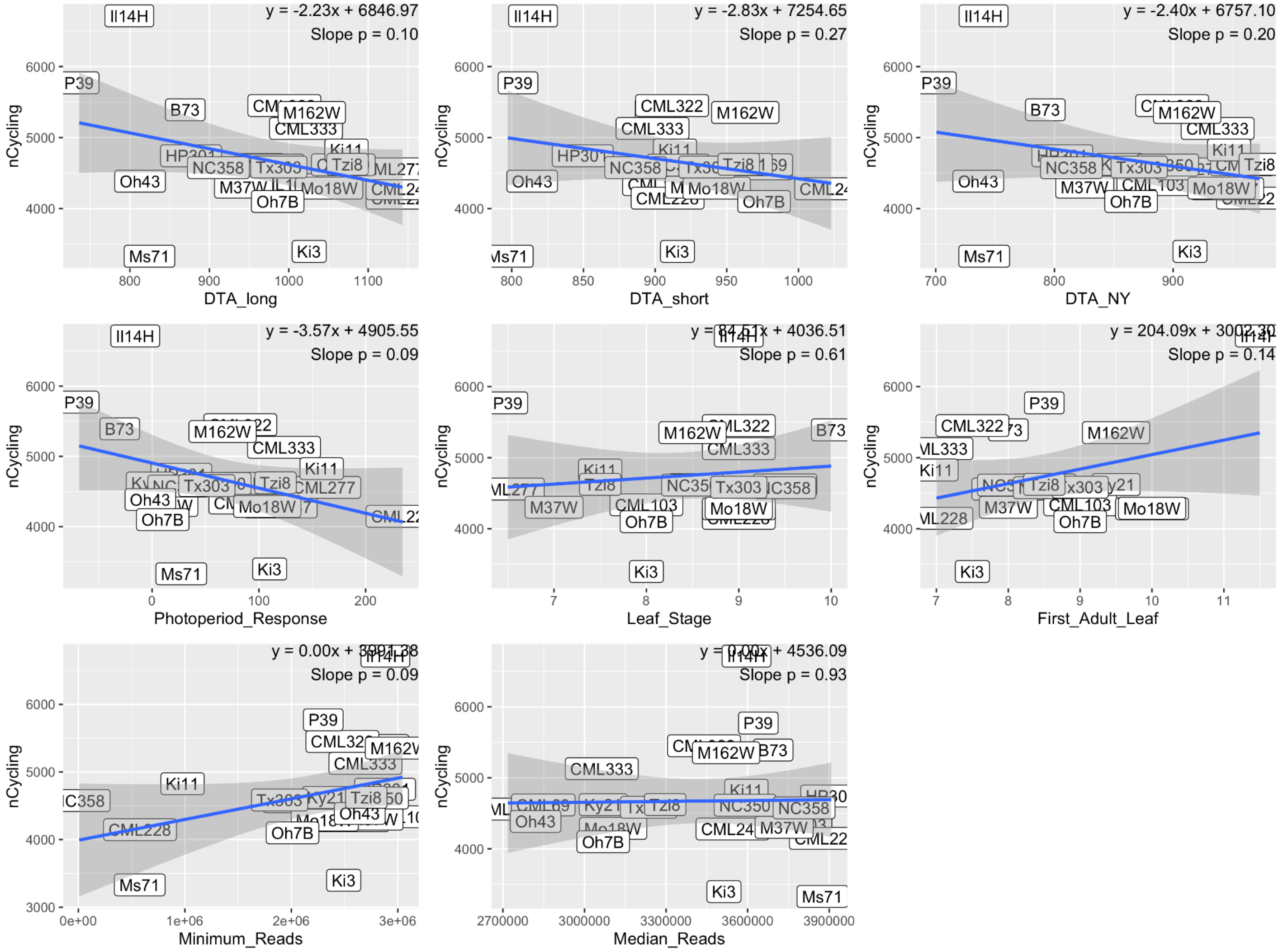

*Supplemental Figure 2: Motif logos from convolutional filters of the CNN trained to predict rhythmicity from promoter sequence*

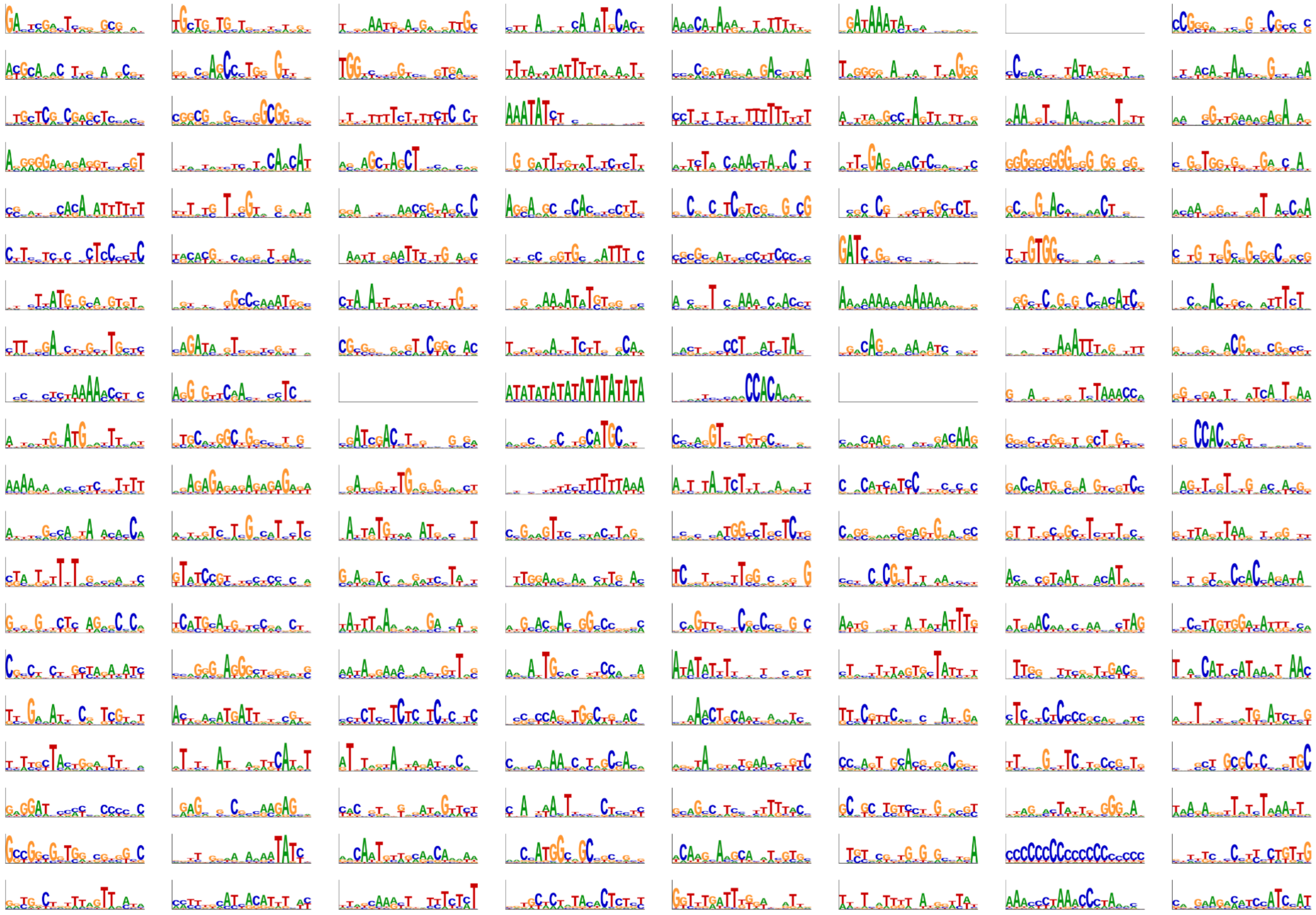

*Supplemental Figure NN: number of genes identified as cycling in three inbreds grown under field conditions and in the growth chamber study.*

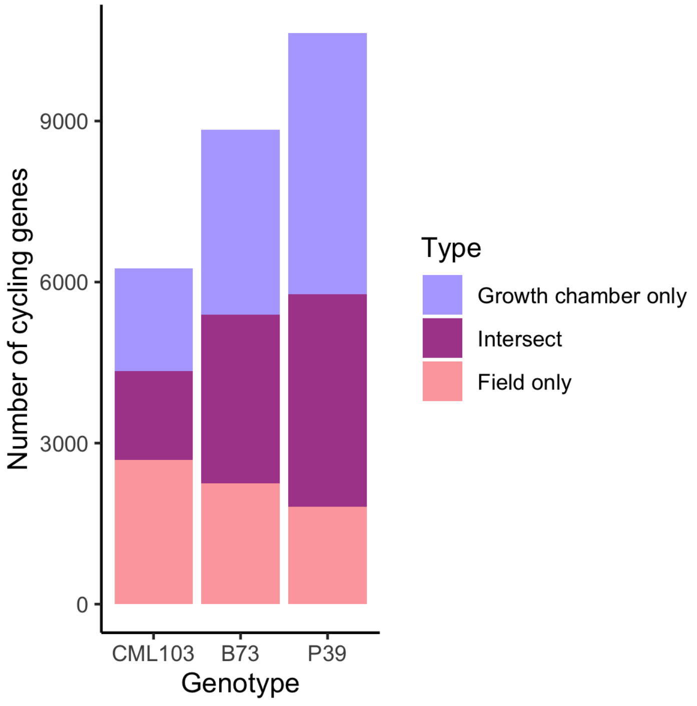
